# Genome size, rather than sociality, predicts the turnover of duplicated genes in termites and hymenopterans

**DOI:** 10.64898/2026.07.20.739705

**Authors:** Cong Liu, Cédric Aumont, Yi-Ming Weng, Alina A. Mikhailova, Aleš Buček, Jan Šobotník, Mark C. Harrison, Dino P. McMahon, Simon Hellemans, Thomas Bourguignon

**Affiliations:** Okinawa Institute of Science & Technology Graduate University, 1919-1 Tancha, Onna-son, 904-0495 Okinawa, Japan; Institute for Evolution and Biodiversity, University of Münster, Hüfferstraße 1, 48149 Münster, Germany; Department for Materials and the Environment, BAM Federal Institute for Materials Research and Testing, 12205, Berlin, Germany; Biology Centre of the Czech Academy of Sciences, Institute of Entomology, České Budějovice, 37001, Czech Republic; Faculty of Tropical AgriSciences, Czech University of Life Sciences, Kamýcká 129, 16521 Prague, Czech Republic; Centre for Discoveries in Life Sciences, Coventry University, Coventry, UK; Institute of Biology, Freie Universität Berlin, 14195, Berlin, Germany

## Abstract

Gene duplication is a major source of genetic variation and is considered as an important driver of evolutionary novelties, including eusociality in insects, such as ants, wasps, bees, and termites. However, it remains unclear whether selection acts to increase gene copy number in social insects. Here, we studied the paranomes, namely, the whole set of paralogous genes in a genome, of Blattodea and Hymenoptera. We estimated the rates of gene duplication and loss using the distribution of synonymous substitution rate of paralogous genes and showed that regardless of sociality, duplicated genes were lost more rapidly than expected under random drift, indicating that negative selection on duplicated genes is prevalent across Blattodea and Hymenoptera. The rates of gene duplication and loss varied independently of sociality levels, but were positively related to genome size, suggesting the expansion/contraction of gene families can be a side effect of genome expansion/contraction. These results call for a reevaluation of adaptative gene duplications.

## Introduction

Gene duplication is a major source of new genes and functions (Ohno 1970; Cisneros et al. 2026), contributing to evolutionary innovations and adaptation to new environments (Kondrashov 2012). For example, gene duplication was proposed to have contributed to the origin of eusociality (Gadagkar 1997), a major evolutionary transition involving a reproductive division of labor amongst related individuals and the emergence of castes (Szathmáry and Smith 1995; Wilson and Hölldobler 2005; Nowak et al. 2010). Recent comparative transcriptomic investigations revealed caste-biased expression for hundreds of paralogous genes in bees (Chau and Goodisman 2017; Xu and Colgan 2025) and termites (Shigenobu et al. 2022; Fujiwara et al. 2025), which were interpreted as supporting a prominent role of gene duplications in the evolution of eusociality in insects. However, the identity of genes differentially expressed among castes is largely inconsistent amongst species (Harrison et al. 2021), casting doubt on the contribution of gene duplication to the emergence of castes in the ancestor of social insects and, more generally, to insect social evolution.

To investigate the adaptive role of gene duplication in the evolution of eusociality, it is crucial to determine the rate at which duplicated genes are generated and lost across insect species with various sociality levels. However, limited information is available on the rate of gene duplications. Direct measures of spontaneous gene duplication rates are only available for a few model organisms and indicate rates vary between 10^-5^ and 10^-7^ duplications per gene per generation (Lynch et al. 2008; Schrider et al. 2013; Konrad et al. 2018; Chain et al. 2019; Gill et al. 2023). Therefore, gene duplications are rare events, difficult to measure through parent-offspring comparison (Wang et al. 2023). An alternative approach is to estimate the rates of gene duplication and loss from multiple genomes with known divergence time, by counting such events on the branches of a species phylogenetic tree (e.g., Gao and Innan 2004; Zwaenepoel and Van de Peer 2019), as is often done to infer the single-nucleotide mutation rate (Kimura 1968; Bergeron et al. 2023). Measures of gene duplication rate on tree branches suggested a higher duplication rate in honeybees than in several related species with lower levels of sociality (Chau and Goodisman 2017). However, the results were not significant after correction for phylogenetic non-independence. In addition, assuming a constant gene duplication rate (*i.e*., the number of newly introduced gene lineages per gene per time unit), the number of gene duplication events accumulates exponentially over time; therefore, counting gene duplication events over an arbitrary interval of time does not provide an estimation of gene duplication rate. Consequently, the selective pressure driving the dynamics of duplicated genes across species and its contribution to social evolution remain unclear.

An alternative way to estimate gene duplication rate is to infer it from the age distribution of duplicated genes present in the genomes of modern species. The number of synonymous substitutions per site (*dS*) accumulated after gene duplication provides a proxy for the age of gene duplication events (Vanneste et al. 2013; Li et al. 2018; Zwaenepoel and Van de Peer 2019). During a very short period before the present, gene loss is negligible, and gene duplication events represent the spontaneous rate of gene duplication (Lynch and Conery 2000). The shape of the age distribution of duplicated genes is determined by the rates of gene duplication and loss. If selection acts neutrally on duplicated genes, duplicated genes are retained at the spontaneous gene duplication rate following random genetic drift (Kimura and Ohta, 1971). If selection favors the removal or retention of duplicated genes, gene loss is faster or slower than expected from random genetic drift. Therefore, from the age distribution of gene duplication, it is possible to model the rate of gene duplication and loss and determine the nature of the selective processes acting on duplicated gene copies.

Here, we characterized the paranomes (*i.e*., the entire set of paralogous genes in a genome) of 47 Blattodea (45 termites and two cockroaches) and 41 Hymenoptera and estimated the rates of gene duplication and loss for each species by modeling the *dS* distribution of paralogous genes. We found that gene loss was more frequent than expected by random genetic drift in most species, suggesting that selection favors the removal of duplicated gene copies. The rates of gene duplication and loss did not differ significantly between termites with different levels of sociality and between social and solitary hymenopterans. However, species with large genomes harbored a significantly more abundant and diverse set of paralogous genes and experienced faster turnover of duplicated genes than their relatives with small genomes.

## Results and Discussion

### The paranome size and richness are coupled with genome size rather than sociality

We analyzed high-quality genome datasets of two insect lineages, Blattodea (Table S1) and Hymenoptera (Table S2). Both lineages include species with varying levels of sociality. The Blattodea dataset contained the genomes of two cockroaches, *Blatta orientalis* and *Cryptocercus meridianus*, and 45 termite species (Liu et al. 2025) (Table S1). The cockroaches were excluded from phylogenetic comparative tests on the correlations with sociality. The 45 termite species were classified into two groups based on their level of sociality, which is best approximated by their ontogeny. Species with a linear ontogeny are considered to have low levels of sociality. They have old immatures called pseudergates that perform the colony work while retaining the capacity to develop into alate imagoes (Roisin 2021; Roisin and Korb 2010; Mizumoto and Bourguignon 2021). In contrast, species with a bifurcated ontogeny are considered to possess a high level of sociality. They have a true worker caste that irreversibly deviates from the alate developmental line at an early stage of development (Roisin and Korb 2010). The group of highly social species contained 30 species forming three lineages: (1) Mastotermitidae, (2) Rhinotermitidae, and (3) Heterotermitidae + Termitidae (Geoisoptera). The other group, which contained species with lower levels of sociality, contained 15 species. All 47 genomes were assembled from long reads, and 22 were scaffolded to a near-chromosomal level (Liu et al. 2025).

The Hymenoptera dataset contained genomes from 41 species available on InsectBase 2.0 (Mei et al. 2022), including 13 solitary species and 28 social species. The social species formed three lineages that independently acquired eusociality (Branstetter et al. 2017; Peters et al. 2017; Piekarski et al. 2018; Bossert et al. 2019): (1) ants, (2) paper wasps (Polistinae) and true wasps (Vespinae), and (3) bees, including *Apis* honeybees, *Bombus* bumblebees, and *Melipona* stingless bees (Table S2). All the hymenopteran genomes were scaffolded, with 33 at a near chromosomal level.

Using the two genome datasets, we independently compared the paranomes of (1) termites with linear ontogeny and bifurcated ontogeny and (2) social and solitary hymenopterans. We did not directly compare termites and hymenopterans because the two lineages diverged nearly 400 million years ago (TimeTree database, Kumar et al. 2022) and have evolved distinct biology and ecology. We first clustered genes into hierarchical orthologous groups (HOGs) using OrthoFinder (Emms and Kelly 2019) for each dataset separately and extracted multiple-copy HOGs to estimate paranome characteristics for each species (Table S1, S2). Genomes of Blattodea contained between 11,286 and 15,148 HOGs, including 584 to 3262 multiple-copy HOGs (Figure S1A, S1C). The paranome size, expressed as the total number of genes contained in multiple-copy HOGs, varied between 1538 and 9802 genes per genome in Blattodea (Figure S1A, S1B). The number of HOGs was more variable in Hymenoptera. Each hymenopteran genome contained between 8,828 and 22,543 HOGs. The paranome richness ranged from 581 to 3781 multiple-copy HOGs, and the paranome size varied between 581 and 23,751 genes (Figure S1D-F).

The expansion of the paranome size may be linked to the acquisition of new traits (Ohno 1970; Magadum et al. 2013; Wapinski et al. 2007; Vollger et al. 2022), such as the evolution of castes in social insects (Gadagkar 1997). It could also be a side effect of genome expansion and contraction, as large genomes tend to have more genes (Hou and Lin 2009; Friar et al. 2012; Elliott and Gregory 2015; Liu et al. 2025). To investigate the expansions and contractions of the paranome size, we performed phylogenetic generalized least squares (PGLS) (Table S4, Figure 1) and found that genome size significantly correlated with paranome size and richness in both termites (Figure 1A-B) and Hymenoptera (excluding *Xylocopa violacea* as an outlier with extraordinarily large genome) (Figure 1C-D) (*P* < 0.006, R^2^ > 0.47). In contrast, levels of sociality did not significantly correlate with paranome size and richness (*P* > 0.58) (Figure 1). The absence of a correlation between the paranome size and richness and sociality was further supported by simulation-based phylogenetic analysis of variance (pANOVA) (*P* > 0.37) (Table S5), while the effect of genome size was confirmed using a Spearman’s correlation test performed on phylogenetic independent contrasts (PICs) (*P* < 0.023, Rho > 0.36) (Table S6). Therefore, our results show that the expansion and contraction of social insect paranomes are linked to variation in genome size but not to variations in levels of sociality.

**Figure 1.**
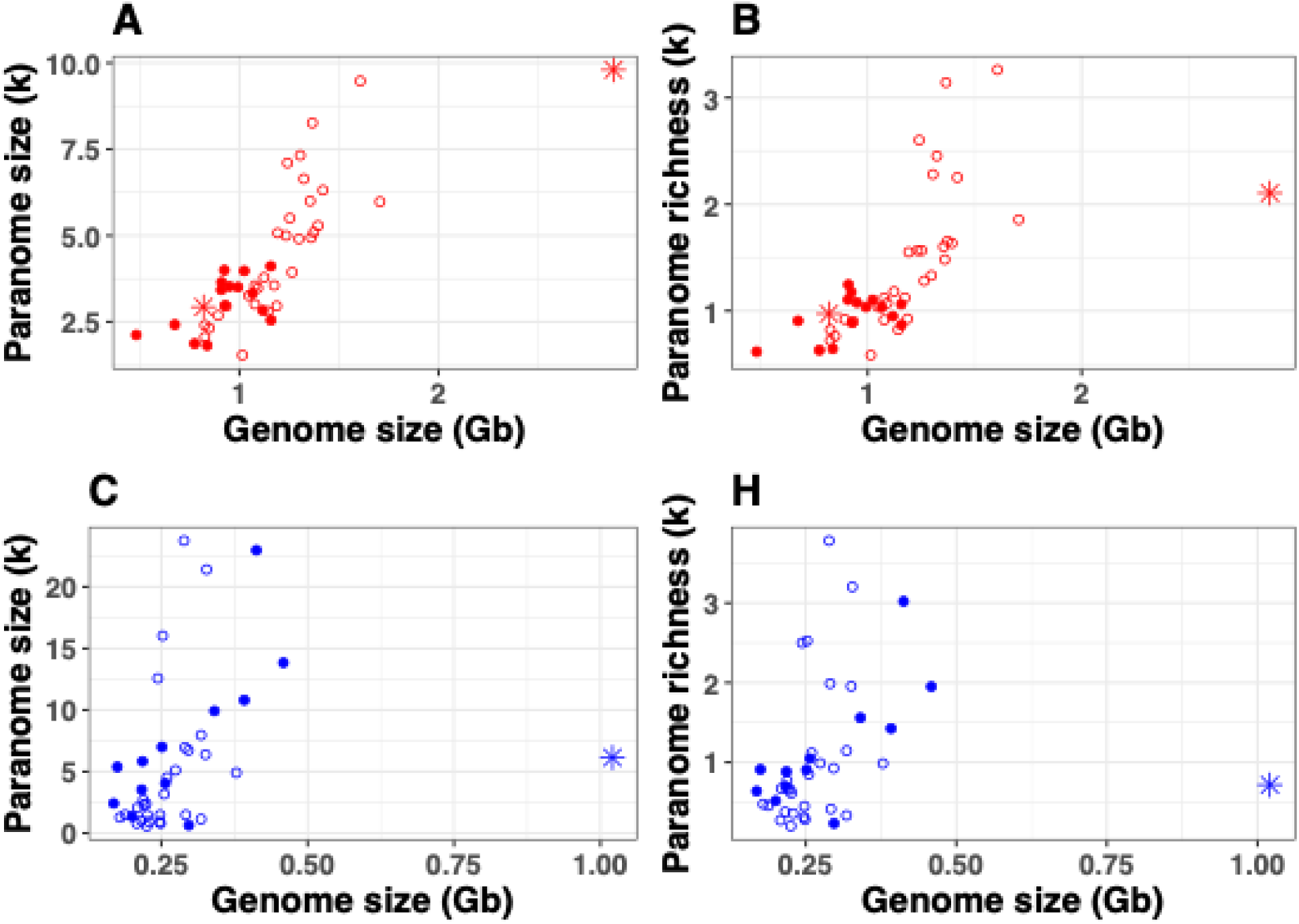
Number of paralogous genes and number of hierarchical orthologous groups (HOGs) containing paralogous genes against genome size for Blattodea (A-B) and Hymenoptera (C-D). The cockroaches (*Blatta orientalis* and *Cryptocercus meridianus*) and the solitary carpenter bee *Xylocopa violacea* were represented by red and blue star points, respectively. Filled red and blue circles represented blattodean and hymenopteran species of higher social complexity, respectively (bifurcated ontogeny for Blattodea; eusociality for Hymenoptera).

### Paranomes expand and contract via small-scale gene duplications and losses rather than large-scale events

The spatial scales of gene duplication events can range from a single gene to the entire genome (Ohno 1970). We searched for gene duplications of large spatial scale through gene synteny analyses of 22 blattodean and 33 hymenopteran genomes assembled at a near-chromosome level. Using reciprocal homology searches, we identified and compared intra- and inter-genome synteny blocks containing five or more genes. Overall, our results revealed (i) fewer intra-genome synteny blocks (Blattodea, mean: 164, median: 150.5; Hymenoptera, mean: 515, median: 227) than inter-genome blocks (Blattodea: mean: 420.36, median: 417; Hymenoptera, mean: 589.74, median: 604) (one-sided Wilcoxon tests, Blattodea: *P* = 3.14e-14, Hymenoptera: *P* = 1.36e-07, Figure S2A); (ii) shorter intra-genome synteny blocks (Blattodea, mean: 7.96, median: 6; Hymenoptera, mean: 7.57, median: 6) than inter-genome blocks (Blattodea, mean: 16.41, median: 8; Hymenoptera, mean: 10.32, median: 7) (one-side Wilcoxon tests, *P* < 2.2e-16, Figure S2B); (iii) a smaller proportion of genes being part of intra-genome blocks (on average, 14.88% and 30.53% of all genes in Blattodea and Hymenoptera, respectively) than inter-genome blocks (40.30% for Blattodea and 31.25% for Hymenoptera on average) (one-side Wilcoxon tests, Blattodea: *P* = 9.25e-15, Hymenoptera: *P* = 0.033; Figure S2C). Altogether, these results indicate that intra-genome synteny blocks were few and fragmentary, supporting the absence of large segmental duplications in blattodean and hymenopteran genomes because such events introduce large intra-genome synteny blocks. Furthermore, we found a significant linear correlation between the proportion of genes in inter-genome blocks and the divergence of fourfold degenerated sites (4dtv) between species pairs in both Blattodea and Hymenoptera (*P* < 2e-16; Blattodea: R^2^ = 0.67, Hymenoptera: R^2^ = 0.41; Figure S2D-E), suggesting a gradual loss of inter-genome gene synteny as species diverge. These results refute the scenario that extensive genome scrambling following segmental duplications eliminated intra-genome synteny blocks. Therefore, the paranomes of Blattodea and Hymenoptera have been continuously expanding primarily through small-scale gene duplications rather than large segmental duplications.

### Duplicated genes are negatively selected across insects with various levels of sociality

We modeled the process of gene duplication and loss that generated the paranomes encoded in the genomes of modern species. We estimated the net gene duplication rate from the age distribution of duplicated genes retained in modern genomes (Equation 7), using *dS* as a proxy for the age of duplication events. If selection on duplicated genes is neutral, the net gene duplication rate equals the spontaneous gene duplication rate A (Kimura and Ohta, 1971) (the constant-rate model, Equation 9). If selection acts to remove or retain duplicated genes, gene loss is higher or lower than expected from the constant-rate model and the net gene duplication rate decreases or increases as we go backward in time. For these latter scenarios, we assumed the net gene duplication rate changes exponentially over time, with its monotonicity determined by the parameter a (Equation 8), and the age distribution given by the decreasing-rate (Equation 10) and increasing-rate models (Equation 11), respectively.

We built the *dS* distributions of paralogous genes of Blattodea and Hymenoptera genomes and fitted the three models (Equations 9-11) separately. Model selection was carried out using the Bayesian information criterion (BIC) (Figure 2, S3, S4). The distributions were all approximately L-shaped, with most gene duplication events being recent, suggesting no apparent burst of gene duplications at any point in time during the evolution of Blattodea and Hymenoptera.

**Figure 2.**
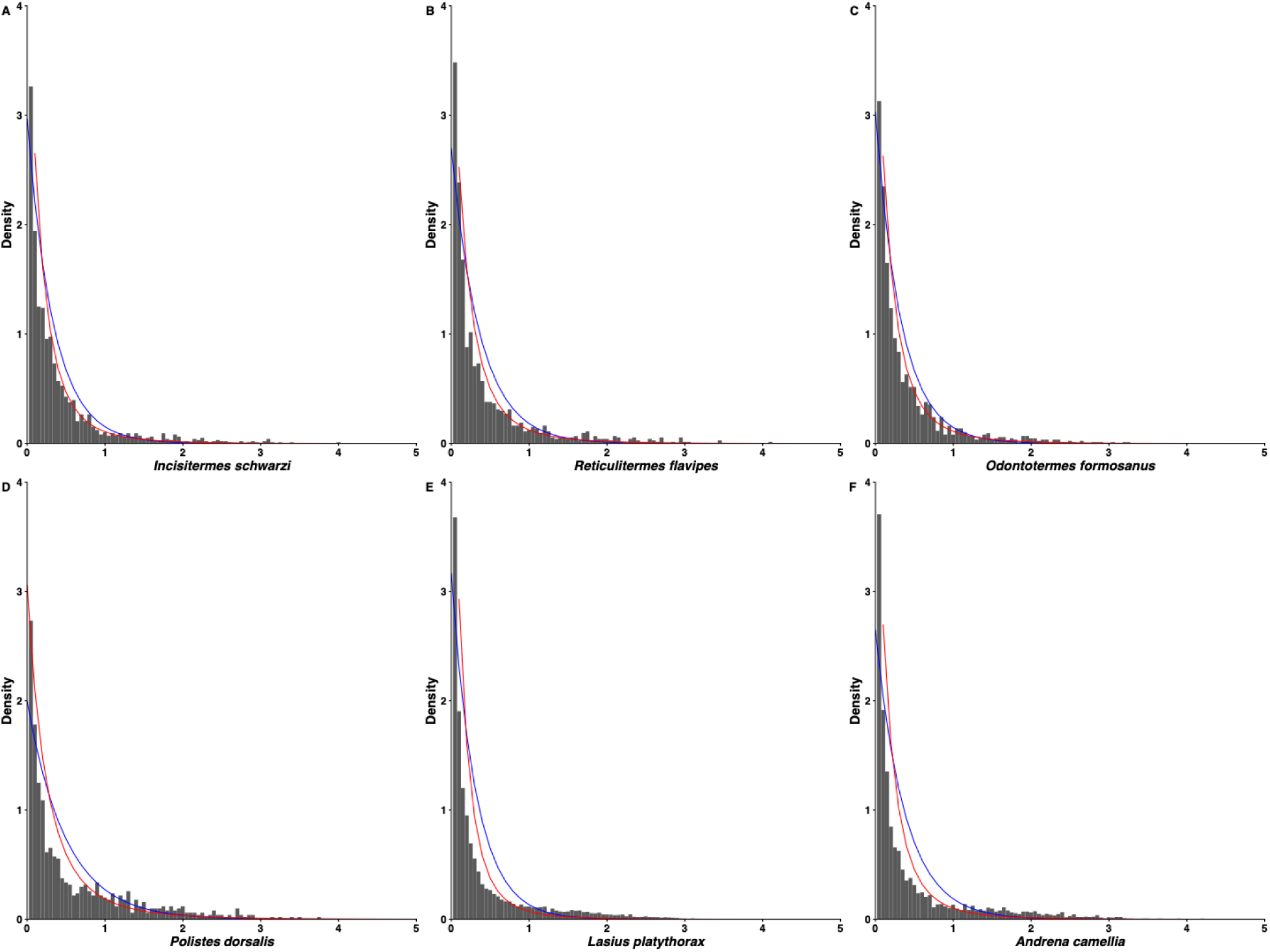
Examples of the synonymous substitution rate (*dS*) distribution of gene duplication events from *Blattodea* (A-C) and *Hymenoptera* (D-F). The blue line is the fitting of the constant-rate model (Equation 9), and the red line is the fitting of the decreasing-rate model (Equation 10).

In all 47 Blattodea species, the decreasing-rate model (Equation 10) was preferred, with BIC weight (BICw) above 99.9% (Figure S1, Table S1), indicating duplicated genes are lost at a faster rate than expected under the constant-rate model (Equation 9). The coefficient λ ranged between 1.969 and 5.641 (mean: 3.805, median: 3.838) and the coefficient a between 1.049 and 1.805 (mean: 1.417, median: 1.398). PGLS showed that both coefficients were not significantly correlated with the levels of sociality inferred from ontogeny (*P* > 0.75), and the correlation with genome size was marginally significant for λ (*P* = 0.063), but insignificant for a (*P* = 0.18) (Figure 3A-B, Table S4). Additionally, pANOVA did not detect an effect of sociality on either coefficient (*P* > 0.91, Table S5). However, a Spearman’s correlation test performed on PICs revealed a significant correlation between genome size and λ (*P* = 0.045, Rho = 0.30), but not between genome size and a (*P* = 0.19, Rho = 0.20) (Table S6).

**Figure 3.**
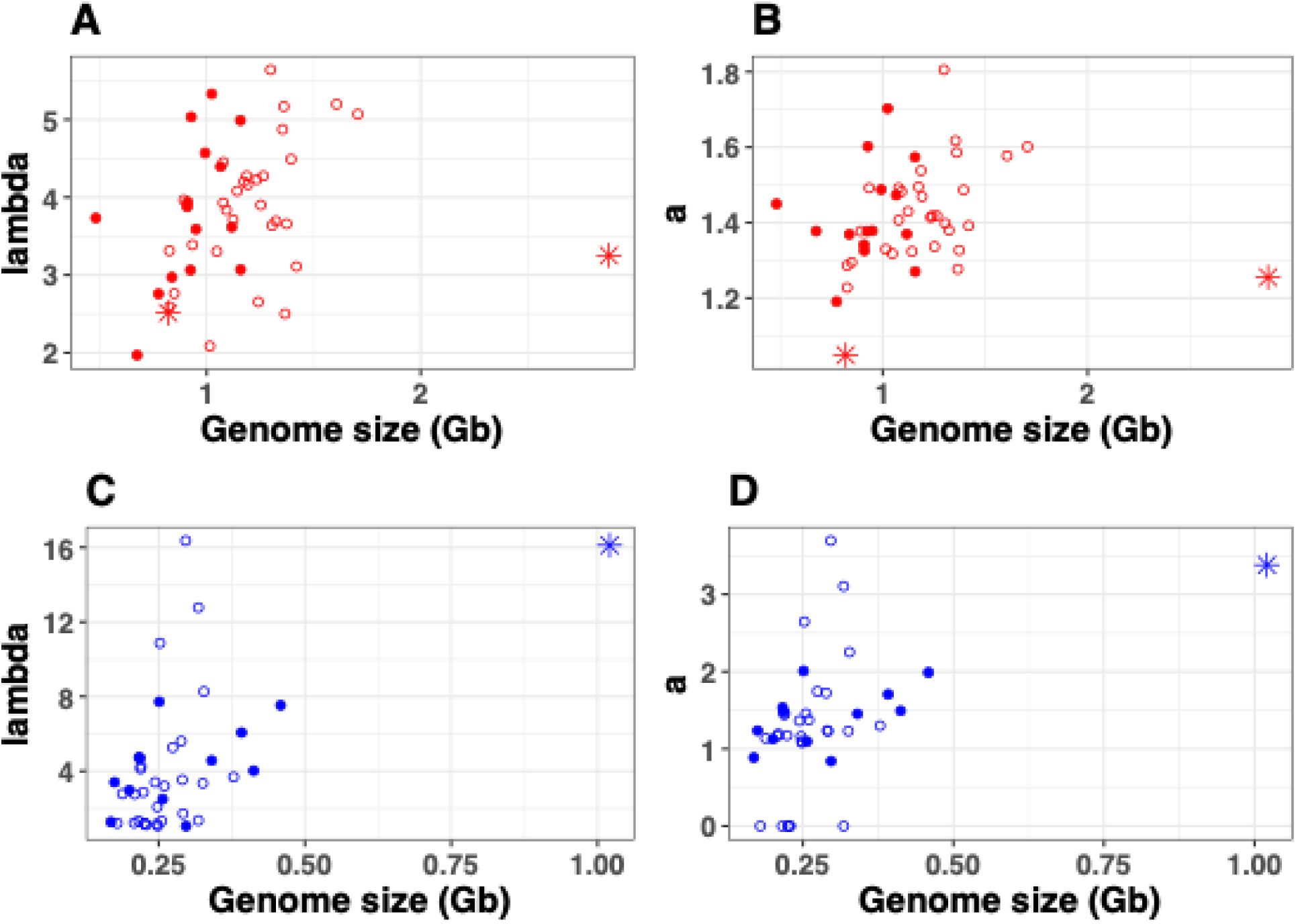
Relationships between genome size and two model parameters: the coefficient A, which quantifies the spontaneous gene duplication rate, and the coefficient *a*, which quantifies selective pressure removing duplicated genes. A-B: Blattodea. C-D: *Hymenoptera*. The cockroaches and solitary carpenter bee *Xylocopa violacea* were represented by red and blue star points, respectively. Filled circles represented species of higher social complexity (bifurcated ontogeny for Blattodea; eusociality for Hymenoptera).

In Hymenoptera (Figure S2, Table S2), the decreasing-rate model had a BICw above 99% in 32 species, and was 90.3% in *Polistes dominula*, 85.3% in *Bombus impatiens*, and 63.1% in *Bombus terrestris*. The constant-rate model was preferred for two honeybees, *Apis cerana* and *Apis laboriosa*, with BICw values of 80.6% and 93.7%, respectively. For the remaining four species (*Apis florea*, *Apis mellifera*, *Atta cephalotes*, and *Vespula pensylvanica*), only the constant-rate model could be fitted. Therefore, the constant-rate models (*a*= 0) were preferred for six species of Hymenoptera, while the best-fit model for the remaining 35 species was the decreasing-rate model (*a*< 0). The coefficient λ ranged from 1.050 to 16.360 (mean: 4.28, median: 3.34), and the coefficient a from 0 to 3.686 (mean: 1.35, median: 1.24). PGLS did not detect a correlation between either model coefficients and sociality (*P* > 0.94) but identified a significant correlation with genome size (λ: *P*=0.0022; a: *P*=0.0067) (Table S4, Figure 3C-D). The non-significant effect of sociality was further confirmed by pANOVA (*P* > 0.72, Table S5), and the significant effect of genome size was corroborated by a Spearman’s correlation test on PICs (λ: *P*=0.0022, Rho = 0.48; a: *P*=0.0039, Rho = 0.45) (Table S6).

Overall, these results showed that the characteristics of the paranome, including its size, richness, and the turnover rate of paralogous genes, do not significantly correlate with sociality levels, and that selective pressures acting against duplicated genes are prevalent across Blattodea and Hymenoptera presenting various levels of sociality. This negative selection against duplicated gene copies at macroevolutionary timescales is in line with the deleterious effects of gene duplications at microevolutionary scales (Zhang et al. 2021). Gene duplication can be deleterious through various mechanisms, such as dosage effects and competition for regulatory elements (Rice and McLysaght 2017; Marques et al. 2011). While the general trend of negative selection on gene duplication is clear, the effect likely varies depending on the functions of duplicated genes. For example, the duplication of universally single-copy genes (Simão et al. 2015) is likely to be highly deleterious and rapidly removed by selection, while some gene families have experienced dramatic expansion in specific lineages, such as the odorant receptors in several lineages of Blattodea and Hymenoptera (Robertson and Wanner 2006; Legan et al. 2021; Gautam et al. 2024; Mikhailova et al. 2025). Whether these gene family expansions are driven by positive selection on gene copies and linked to the evolution of sociality is debated (Gautam et al. 2024; Pellen et al. 2026).

Our results highlight the correlation between genome size and paranome characteristics. Species with larger genomes tend to have more abundant and diversified sets of paralogous genes and undergo a more rapid turnover of paralogous genes. Since protein-coding genes generally account for a very small proportion of insect genomes, gene duplication and loss are unlikely to drive the variations in genome size. On the contrary, processes that impact genome expansion and contraction can, as a side effect, affect the turnover of paralogous genes. For example, genome-scale decrease in DNA loss can promote the retention of duplicated genes. The activity of transposable elements (TEs) (Elliott and Gregory 2025; Canapa et al. 2015; Petersen et al. 2019) can mediate the gain and loss of paralogous genes via processes such as unequal crossing-over between TE insertions and retrotranspositions. Therefore, the expansion and contraction of gene families do not necessarily imply that natural selection favours or discards the retention of duplicated gene copies but may represent a side effect of genome size changes. For example, previous genome analyses performed on Blattodea have repeatedly found simultaneous gene family contractions and genome shrinkage in the ancestor of *Cryptocercus* and termites, and gene family expansion coupled with genome size inflation in the lineage of Termitidae that diverged from Macrotermitinae and Sphaerotermitinae (Mikhailova et al. 2025; Liu et al. 2025; Cui et al. 2026). Our results call for a reappraisal of the criteria used to detect signals of selection on gene family size. Future work aiming at separating the effect of natural selection on gene family expansion/contraction from the general trend of genome expansion/contraction is needed to investigate the adaptive role of gene copy number variations.

## Conclusions

We developed a novel framework for estimating the turnover rate of paralogous genes in a genome and showed rapid gene loss driven by negative selection against gene copies across Blattodea and Hymenoptera. Notably, gene duplication and loss rates were not coupled with sociality, but with genome size, as species with larger genomes experience an accelerated turnover of duplicated genes. While positive selection on specific gene families may have contributed to the evolution of eusociality, gene duplication remains largely deleterious and selected against across social insects. Future development of methods detecting gene family expansions and contractions should integrate the evolutionary dynamics of genome size in order to reliably identify adaptive changes in gene copy number.

## Materials and Methods

### Biological datasets

We used genome data from the two major insect clades containing lineages that independently evolved eusociality: Blattodea and Hymenoptera. The Blattodea dataset contained the annotated genomes published by Liu et al. (2025), including 45 termites, the subsocial woodroach *Cryptocercus meridianus*, and the solitary cockroach *Blatta orientalis* (Table S1). Termites exhibit various levels of social complexity that were assessed based on their ontogeny (Roisin et al. 2011) (Table S1). The Hymenoptera dataset consisted of the genomes of 41 species retrieved from InsectBase 2.0 (Mei et al. 2022), including 13 solitary species and 28 eusocial species (Table S2), spanning three independent origins of sociality (Branstetter et al. 2017; Peters et al. 2017; Piekarski et al. 2018; Bossert et al. 2019): (i) ants, (ii) paper wasps (Polistinae) and true wasps (Vespinae), and (iii) bees, including *Apis honeybees*, *Bombus bumblebees*, and *Melipona stingless* bees. Hymenoptera sociality was classified as solitary or social (Table S2). All analyses were separately performed for Blattodea and Hymenoptera. We compared termites with bifurcated ontogeny to termites with linear ontogeny, which have high and low levels of sociality, respectively. And we compared eusocial hymenopterans with solitary hymenopterans. We did not directly compare termites with hymenopterans.

### Identification of hierarchical orthologous groups

We reconstructed paranomes to determine whether gene duplication is associated with social evolution. The first step was to identify hierarchical orthologous groups (HOGs). For Blattodea, we used the genome annotations of Liu et al. (2025). For Hymenoptera, we used the annotated gene sets available on InsectBase, from which we removed genes containing in-frame stop codons or incomplete reading frames and kept the longest transcript for each gene following the procedure described in Liu et al. (2025).

We reconstructed two species trees, one for Blattodea and one for Hymenoptera, to guide OrthoFinder v.2.5.4 (Emms and Kelly 2019) with the determination of HOGs. We used fourfold degenerated sites (4dtv) of universal single-copy hierarchical orthologous groups (scHOGs) for phylogenetic reconstruction. An initial run of OrthoFinder without a specified species tree identified 1,410 and 1,793 scHOGs for Blattodea and Hymenoptera, respectively. Protein sequences of scHOGs were aligned using MAFFT v.7.508 (Katoh et al. 2002) with the option –*auto*. Protein alignments were converted into codon alignments using PAL2NAL v.14 (Suyama et al. 2006). The 4dtv sites were extracted from codon alignments and used for phylogenetic reconstruction with IQ-TREE v.2.3.6 (Minh et al. 2020). For Blattodea, we fixed the species tree topology to that of the maximum likelihood phylogenetic tree of Liu et al. (2025), reconstructed based on ultraconserved elements, and computed branch lengths using 41,729 4dtv sites. For Hymenoptera, 119,980 4dtv sites were used to reconstruct both the tree topology and branch lengths. A GTR+F+R4 nucleotide substitution model was assigned to both trees as selected by ModelFinder (Kalyaanamoorthy et al. 2017). The resulting Hymenoptera tree topology was congruent with previously published phylogenetic trees (Branstetter et al. 2017; Peters et al. 2017; Sann et al. 2018; Blaimer et al. 2023; Rosa and Melo 2023; Piekarski et al. 2018; Bank et al. 2017; Bossert et al. 2019).

We ran OrthoFinder separately on our Blattodea and Hymenoptera species trees based on 4dtv sites to determine HOGs. Other settings were left on default. Altogether, we identified 34,594 and 61,290 HOGs for Blattodea and Hymenoptera, respectively.

### Gene synteny

We assessed whether paralogous genes originated from duplications of single genes or large segmental duplications by analyzing gene synteny using WGDI v.0.6.5 (Sun et al. 2022). We restricted our analyses to pairwise comparisons of the 22 Blattodea and 33 Hymenoptera genomes considered highly contiguous genome assemblies, with L90 ranging between 5 and 106 (Tables S1-2). We conducted reciprocal homology search between protein sequences of all possible pairs of blattodean and hymenopteran species using DIAMOND v2.1.7.161 (Buchfink et al. 2015) with the options *--evalue 1e-5 --max-target-seqs 20*. WGDI was run on the DIAMOND output for each species pair comparison and gene synteny blocks containing at least five genes were identified at *P*< 0.2. The comparisons between species primarily identified orthologous synteny blocks, while the comparisons of species against themselves identified paralogous synteny blocks. We used the R function *lm* to test for a correlation between the proportion of genes located within synteny blocks in species pairs and the divergence of species pairs on the 4dtv phylogeny.

### Pairwise synonymous substitution rates of paralogous genes present within one genome

The age of gene duplication can be approximated from the synonymous substitution rates per site (*dS*). We estimated dS for each possible combination of pairs of gene sequences part of an HOG and encoded in a single genome. We obtained pairs of sequence alignments from multiple sequence alignments. Multiple sequence alignments from each HOG were first performed on proteins using MAFFT with the –auto option and converted into codon alignments using PAL2NAL for estimation of synonymous (*dS*) and nonsynonymous (*dN*) substitution rates. We estimated *dS* and *dN* for pairs of sequences using the *kaks* function implemented in the R package seqinr (Charif and Lobry 2007) with the parameter rmgap=FALSE. This approach calculates *dS* and dN for pairs of sequences using the method of Li (1993) and the nucleotide substitution model of Kimura (1980). For pairs of sequences sharing little homology or close to genetic saturation, the *dS* values from seqinr are negative or close to 10. We manually corrected these *dS* values to +10.

### Identifying gene duplication events and their timing

We used the *dS* values obtained for pairs of sequences to estimate divergence times between paralogous genes originating from gene duplication events within one genome (Vanneste et al. 2013; Li et al. 2018; Zwaenepoel and Van de Peer 2019). All analyses were independently performed for each genome. We clustered paralogs into dendrograms using the R function *hclust* with *dS* values as a dissimilarity measure. Internal nodes of these dendrograms represent gene duplication events. For each gene duplication event, we computed the median *dS* across all possible gene pairs that diverged from that node (Zhang et al. 2004; Maere et al. 2005; Vanneste et al. 2013; Li et al. 2018; Zwaenepoel and Van de Peer 2019). This approach allowed us to remove redundant pairwise *dS* values for the HOGs containing three or more paralogs. Gene duplication events with *dS* above 5 were discarded.

### Modeling the age distribution of gene duplications

We built three mathematical models to investigate the dynamics of gene duplication during Blattodea and Hymenoptera evolution. Paranomes are the sum of all the new gene lineages that were retained following the process of gene duplication and loss. The rate at which new gene lineages are retained, or the net gene duplication rate, cannot be negative since gene loss is not always observable. Using *dS* as a proxy for time, we denote the timing of gene duplication events by *s* and the number of genes in the paranome by *n*. We can write *n* as a function of s going backward in time as genes coalesce at each duplication event. Denote by *δ*= δ(s) the net gene duplication rate, and for a small increment *△s*, we obtain

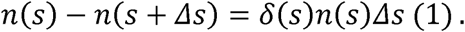

That is,

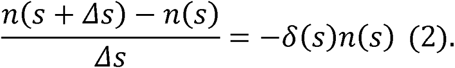

As Δs 0, the Equation 2 can be written as follow

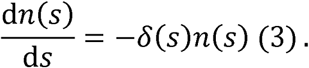

The solution of Equation (3) is

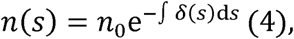

where *n*_δ_ is a constant. Furthermore, the frequency of gene duplication events with time shorter than *s*, that is, the cummulative distribution function of gene duplication events over time, is given by

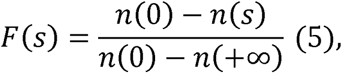

and the corresponding probability density distribution by

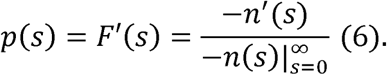

Combining Equation 3 and Equation 6, we can write

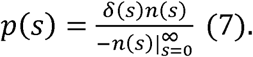

If the duplicated genes are retained following selectively neutral processes, the net gene duplication rate *δ(s)* equals the spontaneous gene duplication rate (Kimura and Ohta, 1971). Otherwise, if there are selective pressures to retain or discard duplicated genes, the net gene duplication rate is expected to change over time. We assume that the net gene duplication rate changed over time exponentially

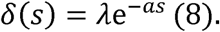

The coefficient λ represents the net gene duplication rate at *s*= 0, serving as a proxy for the spontaneous gene duplication rate. The coefficient a determines the direction and magnitude of change in net gene duplication rate *δ(s)* along s and serves as a proxy for the strength of selection for retaining or discarding duplicated genes.

If *a*= 0, the net gene duplication rate is a constant λ (constant-rate model), suggesting the retention of duplicated genes following random processes. From Equations 4, 7, and 8, the probability density distribution of the timing of gene duplication events *p(s)* follows an exponential decay and can be solved as

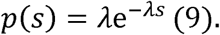

If *a*> 0, the net gene duplication rate decreases along *s* (decreasing-rate model), suggesting that duplicated genes are discarded faster than expected from random retention.

In this scenario, natural selection favors the loss of duplicated genes, and p(s) can be solved from Equations 4, 7, and 8 as

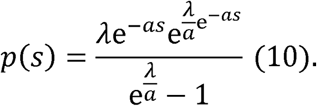

If *a*< 0, *δ(s)* increases along *s* (increasing-rate model), suggesting that duplicated genes are retained at a higher rate than the expectation of the constant-rate model. In this scenario, natural selection acts to retain duplicated genes, and p(s) can be solved from Equations 4, 7, and 8 as

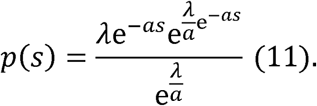

### Fitting the three models of gene duplication events over time

We fitted the three models (Equations 9-11) using the *fitdistr* function implemented in the R package MASS (Venables and Ripley 2002). Models were fitted for each species separately using the combined *dS* values of all gene duplication events. For the constant-rate model (Equation 9), we used the option *densfun=“exponential”* without starting values since the exact maximum-likelihood estimation (MLE) can be calculated analytically. The other two models were fitted through iterations. Since λ represents the net gene duplication rate at *s*= 0, it is largely determined by the probability density around *s*= 0. Therefore, the MLE of the constant-rate model is expected to be a good starting value for the decreasing-rate and increasing-rate models (Equations 10 and 11). We determined the value of the a coefficient in the increasing-rate model by testing the model fitting with a values incrementally increasing by 0.1 from 0.1 to 10. The same approach was used for the decreasing-rate model, but with a values incrementally decreasing by 0.1 within a range varying between −0.1 and −10. Best-fitted a values were selected using the Bayesian information criterion (BIC) with the R function *BIC*. In some cases, the computation did not converge despite a number of iterations higher than *Machine$double.xmax*=1.797693e+308, the maximum number of iterations possible with R. We considered these failures as an indication that the distributions of *dS* values were unlikely to be explained by these models.

For the models fitted successfully, the best-fitted model was selected with the BIC weight (BICw) using the aic.w function implemented in the R package phytools (Liam et al. 2012).

### Statistical analyses

We tested for a correlation between sociality or genome size and the paranome size and richness. We also tested for a correlation between two model parameters (A and *a*) and sociality or genome size. All analyses were performed with comparative phylogenetic methods. We used three methods: phylogenetic generalized least squares (PGLS) (Table S4), simulation-based phylogenetic analysis of variance (pANOVA) (Table S5), and Spearman’s correlation test on phylogenetic independent contrasts (PICs) (Felsenstein 1985) (Table S6). All tests were performed with a time-calibrated species tree. For Blattodea, we used the time-calibrated species tree of Liu et al. (2025). For Hymenoptera, we time-calibrated the species tree reconstructed with the alignment of 119,980 4dtv sites of single-copy HOGs using LSD2 (To et al. 2016) implemented in IQ-TREE. We used 16 calibrations, including the age of the root of the tree (215 million years ago) and the ages of 15 internal nodes obtained from the timetree database (Kumar et al. 2022) (Table S3). A GTR+F+R4 model of nucleotide substitution was assigned, as selected by ModelFinder. The time-calibrated phylogenetic trees of Blattodea and Hymenoptera are provided in Table S7.

We carried out PGLS using the *gls* function implemented in the R package *nlme* (Pinheiro et al. 2023), with the expected covariance computed with the 4dtv tree using the *corBrownian* function implemented in the R package ape (Paradis and Schliep 2019). We performed pANOVA using the *phyloANOVA* function of the R package phytools (Garland et al. 1993, Revell et al. 2012). PIC was computed using the pic function implemented in ape. Spearman’s correlation tests were performed using the R function cor.test. Wilcoxon tests were performed using the R function wilcox.test.

## Author contributions

C.L., S.H., and T.B. conceptualized the experiments. C.L. analysed the data. C.L. and T.B. wrote the manuscript. S.H., A.A.M., C.A., Y.M.W., A.B., J.S., M.C.H., and D.P.M. read and edited the manuscript and accepted the final version.

## Data Availability

The genome assemblies of the 47 Blattodean species used in this study are available on GeneBank under bioproject PRJNA1198669. The genomes of Hymenopteran species are available in InsectBase.

## Acknowledgements

We thank OIST’s Scientific Computation and Data Analysis Section (SCDA) for providing access to the OIST computing cluster.

## Supplementary Material

**Figure S1.** Characteristics of the paranomes across species of Blattodea and Hymenoptera. Species tree of (A) Blattodea and (D) Hymenoptera. Number of single-copy genes (stars) and paralogous genes (circles) in (B) Blattodea and (E) Hymenoptera. Number of hierarchical orthologous groups (HOGs) containing paralogous genes in (C) Blattodea and (F) Hymenoptera.

**Figure S2.** Intra- and inter-genome synteny across the genomes of 22 Blattodea and 33 Hymenoptera. (A) Number of syntenic blocks containing more than five genes. (B) Length of gene syntenic blocks. (C) Proportion of genes in syntenic blocks. Proportion of genes forming inter-genome syntenic blocks in (D) Blattodea and (E) Hymenoptera plotted against the divergence of fourfold degenerated sites between the corresponding species pairs.

**Figure S3.** The distributions of synonymous substitution rate of gene duplication events in Blattodea. Fitted curve of constant-rate model (blue) and decreasing-rate model (red).

**Figure S4.** The distributions of synonymous substitution rate of gene duplication events in Hymenoptera. Fitted curve of constant-rate model (blue) and decreasing-rate model (red).

**Table S1.** Blattodea genomes.

**Table S2.** Hymenoptera genomes.

**Table S3.** Time calibrations used for the Hymenoptera phylogenetic tree, extracted from timetree.org. It was provided to IQ-TREE as option “--date”.

**Table S4.** Phylogenetic generalized least squares

**Table S5.** Simulation-based phylogenetic analysis of variance.

**Table S6.** Spearman’s correlation test using phylogenetic independent contrasts.

**Table S7.** Time-calibrated phylogenetic trees of Blattodea and Hymenoptera.

