## Supplementary figures and images for "Genome size, rather than sociality, predicts the turnover of duplicated genes in termites and hymenopterans"

### Supplementary Figure S1

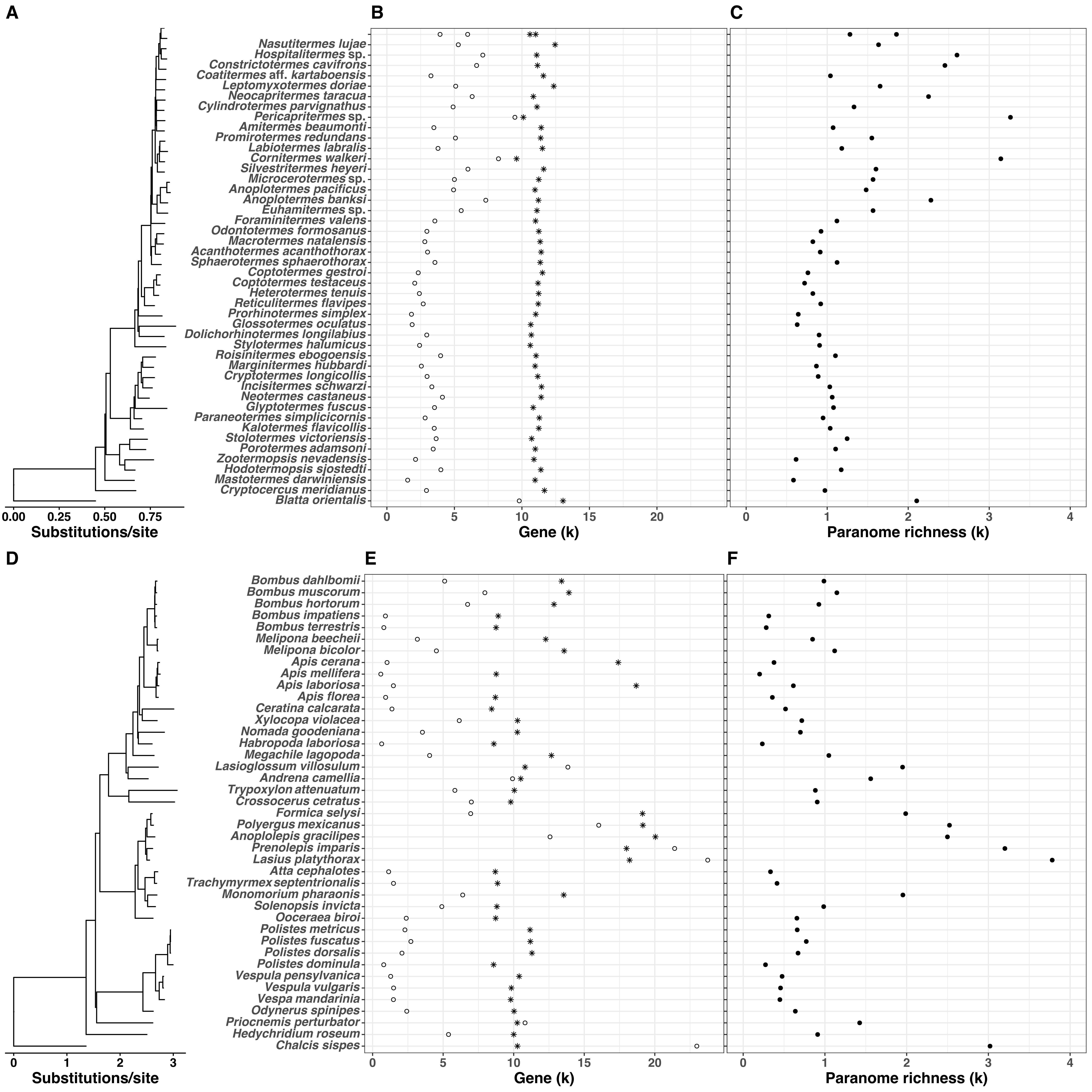

### Supplementary Figure S2

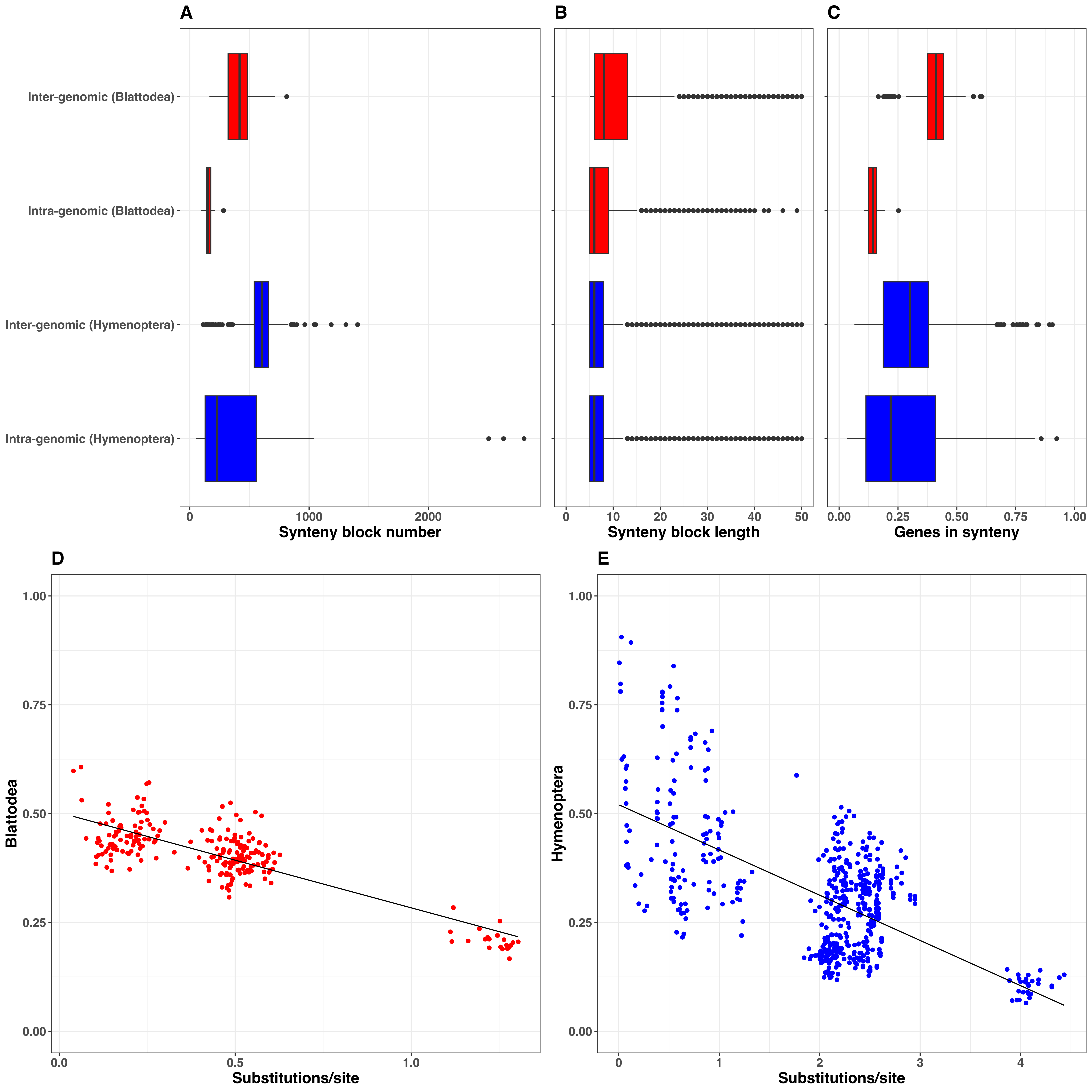

### Supplementary Figure S3

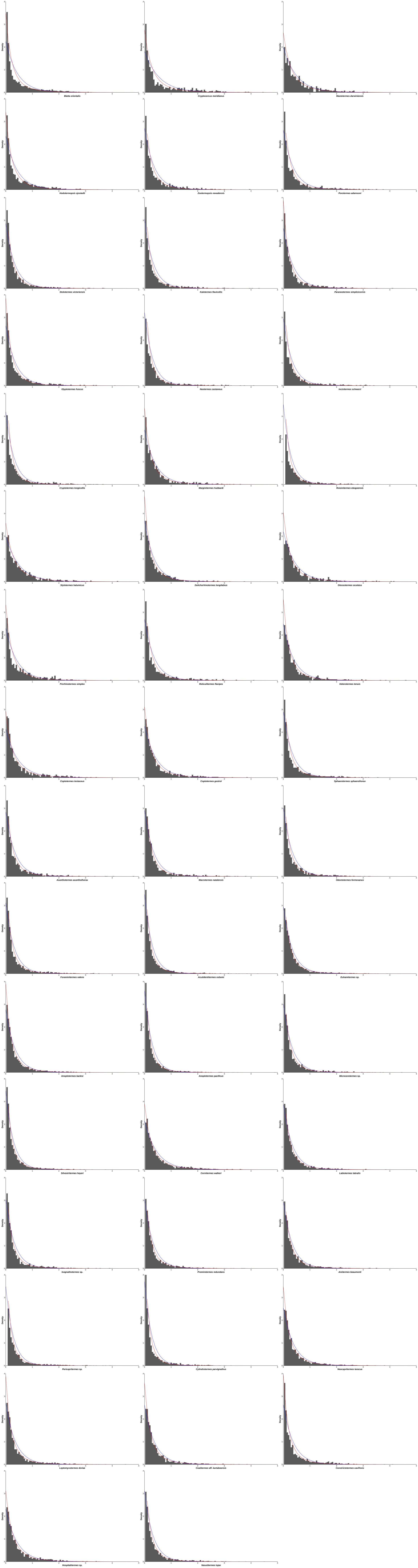

### Supplementary Figure S4

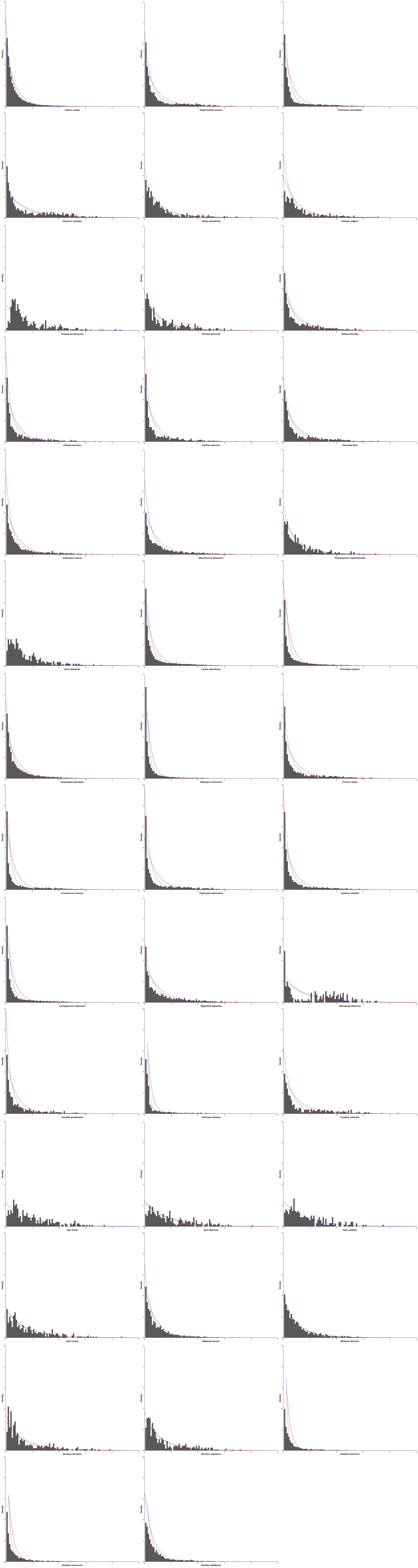
